# Ultra-high field (10.5 T) resting state fMRI in the macaque

**DOI:** 10.1101/2020.05.21.109595

**Authors:** Essa Yacoub, Mark D. Grier, Edward J. Auerbach, Russell L. Lagore, Noam Harel, Kamil Ugurbil, Gregor Adriany, Anna Zilverstand, Benjamin Y. Hayden, Sarah R. Heilbronner, Jan Zimmermann

**Affiliations:** Department of Neuroscience, University of Minnesota, Minneapolis MN 55455; Center for Magnetic Resonance Research, University of Minnesota, Minneapolis MN 55455; Center for Neuroengineering, University of Minnesota, Minneapolis MN 55455; Department of Psychiatry, University of Minnesota, Minneapolis MN 55455; Department of Biomedical Engineering, University of Minnesota, Minneapolis MN 55455; Department of Neurosurgery, University of Minnesota, Minneapolis MN 55455

**Author notes:** Corresponding author: Jan Zimmermann, Department of Neuroscience, Center for Magnetic Resonance Research, University of Minnesota, Minneapolis, MN, 55455.

**Keywords:** Functional connectivity, rhesus macaque, resting-state, spontaneous activity, functional MRI (fMRI)

## Abstract

Resting state functional connectivity refers to the temporal correlations between spontaneous hemodynamic signals obtained using functional magnetic resonance imaging. This technique has demonstrated that the structure and dynamics of identifiable networks are altered in psychiatric and neurological disease states. Thus, resting state network organizations can be used as a diagnostic, or prognostic recovery indicator. However, much about the physiological basis of this technique is unknown. Thus, providing a translational bridge to an optimal animal model, the macaque, in which invasive circuit manipulations are possible, is of utmost importance. Current approaches to resting state measurements in macaques face unique challenges associated with signal-to-noise, the need for invasive contrast agents, and within-subject designs. These limitations can, in principle, be overcome through ultra-high magnetic fields. However, ultra-high field imaging has yet to be adapted for fMRI in macaques. Here, we demonstrate that the combination of high channel count transmitter and receiver arrays, optimized pulse sequences, and careful anesthesia regimens, allows for detailed within-subject resting state analysis at ultra-high resolutions. In this study, we uncover thirty spatially detailed resting state components that are highly robust across individual macaques and closely resemble the quality and findings of connectomes from large human datasets. This detailed map of the rsfMRI ‘macaque connectome’ will be the basis for future neurobiological circuit manipulation work, providing valuable biological insights into human connectomics.

## Introduction

Resting state functional connectivity refers to the temporal correlations between spontaneous blood oxygenation level dependent (BOLD) signals obtained using functional magnetic resonance imaging (fMRI). These fluctuations were first noted in motor cortex (Biswal et al., 1995), but since then many other large-scale networks of correlated temporal patterns in the resting brain have been identified (Biswal et al., 2010; Heuvel and Pol, 2010; Smith et al., 2013; Wang et al., 2009). It was clear even before the advent of fMRI that functional networks at multiple spatial and temporal scales are embedded in the mammalian brain (Essen and Maunsell, 1983; Felleman and Essen, 1991); many years of fMRI neuroimaging have catalogued the spatial and, to some degree, dynamic organization (Allen et al., 2014; Hindriks et al., 2016; Hutchison et al., 2013, 2012) of these networks. These different networks have distinct temporal properties (∼0.01-0.1 Hz fluctuations) and persist through different states such as sleep or anesthesia. Moreover, they are generally consistent across subjects (Smith et al., 2013, 2009) and to some degree generalize across species (Li et al., 2013).

As a method, characterization of resting state networks has proven invaluable for psychiatry, basic neuroscience (Fox and Raichle, 2007; Glasser et al., 2016; Smith et al., 2013), and psychology (Laird et al., 2011). In particular, it offers the ability to simultaneously characterize networks across the whole brain within a short period of time, and without the need to engage the subject in order to collect the data, as opposed to classical task-based experimental designs. These brain networks (and any individual variations) can then be studied across development, aging or disease and provide a high-throughput approach to both basic and clinical human neuroscience.

For several reasons, the resting state technique is a particularly important one to implement in the macaque monkey model. First, macaques offer a critical intermediary between humans and rodent models and have gross neuroanatomy that is highly homologous to that of humans (Hutchison and Everling, 2012; Mars et al., 2016; Petrides and Pandya, 2002; Watson and Platt, 2012). Second, macaques are a valuable model for many complex behaviors and cognitive processes, and thus potentially for psychiatric diseases. Third, macaques offer the opportunity to perform the types of invasive manipulations, including microstimulation, chemogenetics (Galvan et al., 2019; Nagai et al., 2016; Raper et al., 2019; Upright et al., 2018) and optogenetics (Galvan et al., 2017; Khateeb et al., 2019), which we cannot regularity perform on humans. Fourth, additional measurement techniques that are rare or impossible in humans, like single-unit electrophysiological recordings and neuroanatomical tract-tracing, have allowed for careful validation of non-invasive neuroimaging tools such as fMRI and the BOLD response (Logothetis, 2003; Logothetis et al., 2001) or diffusion-weighted-imaging (DWI) (Bakker et al., 2012; Markov et al., 2012). Thus, findings in nonhuman animal models constrain theory and experimentation in human neuroscience, while on the other hand, experimentation using non-invasive tools in humans drives experiments using invasive tools in animals.

Macaques, however, do have a critical disadvantage as a model organism for resting state fMRI. That is, until now, our ability to collect high resolution, high signal-to-noise ratio (SNR) data in macaques has been poor. This is a multifaceted problem resulting mainly from the macaques’ small head size (∼6 % of the human brain volume), the use of standard field strength magnets, the challenges in doing alert studies (especially at high resolutions) and the unavailability of high channel count surface coils. Moreover work on macaques is costly and labor intensive. Thus, sample sizes in classical macaque experiments are generally low (usually 2-4 subjects per experiment), which contradicts the notion of generalization and out-of-sample prediction in human studies. That, in turn, means the utility of neuroimaging in macaques would be greatly enhanced if we could develop ways to improve SNR dramatically. One tool to improve SNR is using monocrystalline iron oxide nanoparticles (MION) as a contrast agent (Moeller et al., 2009; Tsao et al., 2003; Vanduffel et al., 2001). However, this has several disadvantages: (1) the signal then reflects blood volume changes (Leite et al., 2002) rather than the typical BOLD response, complicating its translatability to typical human studies, (2) it has a slower hemodynamic response (Leite et al., 2002; Pelekanos et al., 2020), exacerbating an already major problem with fMRI, and (3) it accumulates in the brain and causes potentially long-lasting health disadvantages. The SNR problem can to some degree also be overcome with larger datasets (i.e. many hours of data collection) but this is expensive and impractical and is accompanied by unavoidable risks to the subjects if general anesthetics are utilized.

One way to overcome the sensitivity problem of acquiring high resolution fMRI is to use ultra-high field strengths in combination with high channel count receiver arrays (Lagore et al., 2020), an approach which has proven extremely productive, to this end, in human studies (Feinberg and Yacoub, 2012; Martino et al., 2018, 2011; Uğurbil et al., 2019, 2003; Yacoub et al., 2001; Zimmermann et al., 2011). While some macaque studies focused on resting state functional magnetic resonance imaging (rsfMRI) (Gilbert et al., 2016; Goulas et al., 2017; Hutchison et al., 2012, 2011; Schaeffer et al., 2018; Vincent et al., 2009, 2007) to date, to our knowledge, resolutions in macaque rsfMRI continue to be greater than 1 mm^3^ as well as longer than 1 second.

Here we explore the advantages of ultra-high field strength neuroimaging for rsfMRI. Using the ultra-high field strength of 10.5 Tesla, combined with a specialized 32 Rx and 8 Tx channel coil, implemented on a human scanner with a commercial console, we demonstrate robust within subject and group ‘connectome’ style resting state fMRI data in 6 lightly anesthetized macaques. Our approach is extremely robust and allows us to repeatedly and robustly measure intrinsic BOLD signals in macaques at high spatial and temporal resolution and with high SNR. Our findings, extending previous approaches, show robust resting state networks similar to human findings that are consistent between macaques and, importantly, identifiable robustly in the individual. Equally important, the components we identify adhere extremely well to anatomical architecture, as well as showing homogeneous sensitivity to cortex and subcortical regions. Our findings demonstrate the general utility that ultra-high field imaging has for in vivo intrinsic BOLD investigations in macaques and non human primates in general.

## Materials and methods

### Animal preparation

We obtained data from 6 adult macaque monkeys (*Macaca fascicularis*: 4 female, and *Macaca mulatta*, 1 male and 1 female). Weights ranged from 3.0 kg to 9.0 kg. Experimental procedures were carried out in accordance with University of Minnesota Institutional Animal Care and Use Committee approval and in accord with the National Institute of Health standards for the care and use of non human primates. All subjects were fed *ad libitum* and pair housed within a light and temperature controlled colony room. Animals were not water restricted. None of the subjects had any prior implant or cranial surgery.

On scanning days, anesthesia was first induced by intramuscular injection of atropine (0.5 mg/kg) and ketamine hydrochloride (7.5 mg/kg). Subjects were transported to the scanner anteroom and intubated using an endotracheal tube. Initial anesthesia was maintained using 1.5-2% isoflurane mixed with oxygen.

Subjects were placed onto a custom-built coil bed with integrated head fixation by placing stereotactic ear bars into the ear canals. For functional imaging, the isoflurane level was lowered to 1%. For *Macaca fascicularis* subjects, an initial bolus injection of 1.5ug/kg fentanyl was administered IV following a continuous administration of 3ug/kg/hr using a syringe pump. Rectal temperature, respiration, end-tidal CO2, electrocardiogram and SpO2 were monitored using an MRI compatible monitor (IRADIMED 3880 MRI Monitor, USA). Temperature was maintained using a circulating water bath as well as chemical heating pads and padding for thermal insulation. Anesthesia was used to eliminate motion effects and physiological stress as well as allow the imaging of surgically naive subjects. Isoflurane is a vasodilator (Farber et al., 1997) and modulates cerebrovascular activity (Vincent et al., 2007). However, previous studies have shown successful synchronous BOLD fluctuations and resting state activity in macaques as well as mice and rats under isoflurane (Hutchison et al., 2012, 2011; Okada et al., 2018; Vincent et al., 2009, 2007).

### Data acquisition

All data were acquired on a passively shielded 10.5 Tesla 60 cm horizontal bore Siemens (Erlangen, Germany) scanner (Magnetom 10.5T Plus) with an SC72 gradient sub-system (Siemens, Erlangen, Germany) operating at a slew rate of 200 mT/m/s. The 10.5T system operates on the E-line (E12U) platform, comparable to clinical platforms (3T Prisma/Skyra, 7T Terra). As such, the user interface and pulse sequences were identical to those running on clinical platforms. A custom in-house built and designed RF coil with an 8-channel transmit/receive end-loaded dipole array of individually 18 cm length combined with a close-fitting 16-channel loop receive array head cap, and an 8-channel loop receive array of 50×100mm size located under the chin (Lagore et al., 2020). The size of 14 individual receive loops of the head cap was 37 mm with 2 larger ear loops of 80 mm - all receiver loops were arranged in an overlapping configuration for nearest neighbor decoupling. The resulting combined 32 receive channels were used for all experiments and supported 3-fold acceleration in phase encoding direction. The coil holder was designed to be a semi-stereotaxic instrument holding the head of the animal in a centered position via customized ear bars. The receive elements were modelled to adhere as close to the surface of the animal’s skulls as possible. Transmit phases were fine-tuned for excitation uniformity for one representative mid-sized animal and the calculated phases were then used for all subsequent acquisitions. Magnetic field homogenization (B0 shimming) was performed using a customized field of view with the Siemens internal 3D mapping routines. Multiple iterations of the shims (using the cardiac adjusted FOV shim parameters) were performed and manual fine adjustments performed on each animal. High order shim elements were disregarded for these procedures.

In all animals a B1 (transmit) fieldmap was acquired using a vendor provided flip angle mapping sequence and then power calibrated for each individual. Following B1 transmit calibration, 3-5 averages (23 minutes) of a T1 weighted magnetization prepared rapid acquisition gradient echo protocol (3D MP-RAGE) were acquired for anatomical processing (TR = 3300 ms, TE = 3.56 ms, TI = 1140 ms, flip angle = 5°, slices = 256, matrix = 320 × 260, acquisition voxel size = 0.5 × 0.5 × 0.5 mm). Images were acquired using in plane acceleration GRAPPA = 2. A resolution and FOV matched T2 weighted 3D turbo spin echo sequence (variable flip angle) was run to facilitate B1 inhomogeneity correction.

Before the start of the functional data acquisition, five images were acquired in both phase encoding directions (R/L, L/R or F/H, H/F) for offline EPI distortion correction. For four monkeys, six runs of 700 continuous 2D multi band EPI (Auerbach et al., 2013; Feinberg et al., 2010; Moeller et al., 2010) functional volumes (TR = 1110 ms; TE = 17.6 ms; flip angle = 60°, slices = 58, matrix = 108 × 154; FOV = 81 × 115.5 mm; acquisition voxel size = 0.75 × 0.75 × 0.75 mm3) were acquired. Images were scanned in left-right phase encoding direction using in plane acceleration factor GRAPPA = 3, partial fourier = 7/8th, and multiband (MB) or simultaneous multislice factor = 2. For the two other monkeys, five runs of 700 continuous 2D multi band EPI functional volumes were acquired (TR = 1200 ms; TE = 17.6 ms; flip angle = 60°, slices = 58, matrix = 106 × 212; FOV = 79.5 × 159 mm; acquisition voxel size = 0.75 × 0.75 × 0.75 mm). Images were scanned in foot-head phase encoding direction using in-plane acceleration factor GRAPPA = 3, partial fourier = 7/8th and MB = 2. The change in sequence parameters was made after upgrading to the 10.5 Plus (E12U) platform which brought with it software changes that made the left right phase encoding more favorable. Since macaques were scanned in sphinx positions, the orientations noted here are what is consistent with a (head first supine) typical human brain study but translate differently to the actual macaque orientation. While left / right orientation is the same, foot-head is actually anterior-posterior in the monkey.

### Image preprocessing

Image processing was performed using a custom pipeline relying on FSL (Jenkinson et al., 2012), ANTs (Avants et al., 2014, 2011), AFNI (Cox, 1996) and a heavily modified CONN (Whitfield-Gabrieli and Nieto-Castanon, 2012) toolbox. Images were first motion corrected using *mcflirt* (registration to the first image). Motion parameters never exceeded 0.5 mm or 0.4° rotation except in one run of one animal, which was discarded (the animal experienced anesthesia induced emesis during the last run). Images were slice time corrected and EPI distortion corrected using *topup*. Ultra-high magnetic fields and large matrix sizes are commonly associated with severe echo planar imaging distortions. Supplementary Fig 1 shows an overview of the extent of the distortion as well as the result of the corrected warps. Anatomical images were nonlinearly warped into the NMT (Seidlitz et al., 2018) template using ANTs and 3DQwarp in AFNI. The distortion correction, motion correction and normalization were performed using a single sinc interpolation. Images were spatially smoothed (FWHM = 2 mm), linear detrended, denoised using a linear regression approach including heart rate, respiration as well as a 5 component nuisance regressor of the masked white matter and cerebrospinal fluid and band-pass filtering (0.008 - 0.09 Hz) (Hallquist et al., 2013). While we chose here to present the data as is typically done in resting state applications, to demonstrate the raw SNR and overall BOLD sensitivity we show an unsmoothed example on the individual macaques brain in supplementary Fig 2.

### Independent component analysis

We used group ICA to uncover ordered reproducible components across subjects. This approach uncovers a set of interpretable group components. Algorithms were used as implemented in the GIFT (Calhoun et al., 2001) software package. Our preprocessing pipeline produced one nuisance corrected concatenated time course of all runs per subject. The temporal dimension of the concatenated time course was then reduced using principal component analysis (PCA). Spatial resting state components were then estimated using the Infomax algorithm approach using the standard GIFT parameters. Time-courses and spatial maps associated with each component were then back projected as implemented in the group-ICA tools in GIFT. Since the process of dimensionality reduction and model selection are somewhat arbitrary (one needs to specify the number of components to estimate), there is no real consensus in how many components to extract. It has been noted that there is no best dimensionality for the underlying neurophysiology of multiple distributed systems (Cole et al., 2010). There are always multiple valid solutions, but we argue that a higher model order (Abou-Elseoud et al., 2009; Kiviniemi et al., 2009; Smith et al., 2009) than previously used (Hutchison et al., 2012, 2011; Vincent et al., 2007) is beneficial in macaques. While one can compartmentalize the brain into networks with associations to visual, motor, sensory, auditory processing, the goal in macaque work is usually shifted towards finer-grain coding to allow for better cross-method comparison. Previous papers have not ignored this but have rather been limited by data quality (e.g., a high dimensional ICA decomposition cannot be obtained from a short rsfMRI run at standard field strength using high resolution). Multiple criteria (Jafri et al., 2008; Li et al., 2007; Zuo et al., 2010) currently exist for optimally selecting the number of independent components for a given dataset. The minimum description length criterion, however, yielded a mean estimation of 703 (SD = 69) independent components for our dataset. To obtain a manageable number of components and to allow for easier comparison with human and previous macaque datasets, 40 components were extracted. To test for the reliability of the decomposition the ICASSO (Li et al., 2007) toolbox was used and ICA iterated 20 times. Lastly, mean group independent components were scaled to empirically derived z-scores. These z-scores are an approximation to the temporal correlation between each voxel and the associated components time-course. A threshold of +/-2 was used as a lower limit of functional connectivity for visualization.

### Identification of Resting State Networks and visualization

Components resulting from the independent component analysis were manually inspected and labeled according to anatomical and functional locations. Since ICA can be extremely noise sensitive, some extracted components are typically associated with motion, scanner noise artifacts or physiological confounds. In our dataset, 6 out of 40 components were discarded as they showed noisy, nonspecific, low correlation activation patterns or correspondence to large veins. Data was both visualized in volume space (custom code written in c++ and openGL) or projected to the surface reconstruction of the NMT template (Seidlitz et al., 2018) using AFNI (Cox, 1996) and SUMA (Saad et al., 2004; Saad and Reynolds, 2012).

### Single subject ICA

Since studies in macaque usually have low numbers of subjects, and because subtle individual features of resting-state connectivity can be lost in group analysis, we also performed single subject ICA. Parameters were matched to those of the group ICA.

### Independent component network connectivity

Temporal correlations between components resulting from spatial ICA analysis as performed here can be high (Calhoun et al., 2003), since our approach to ICA maximizes the statistical independence in space while reducing the dimensionality of the temporal domain. To explore the relationship between the independent components from a temporal perspective we performed functional network connectivity analysis (FNC) (Jafri et al., 2008) between our resulting ICs. Our ICs were further clustered using a purely data driven hierarchical clustering procedure (nonspatial complete-linkage clustering). FNC then analyzes the entire set of connections between pairs of networks in terms of the within- and between-network connectivity sets. Resulting F-statistics were used to test statistically significant connections (connection threshold p<0.05 p-uncorrected, cluster threshold p<0.05 p-FDR corrected using MVPA omnibus test).

## Results

### Group Resting state networks

Group-ICA decomposed the dataset of 6 monkeys into 40 independent components as specified. ICASSO returned a stability index of 0.96 (SD = 0.04) demonstrating that the components are close to orthogonal clusters and highly consistent across multiple ICA iterations. Out of the 40 extracted components, 30 were deemed physiologically relevant, containing more than one anatomical area, adhering to the gray matter as well as displaying a reasonable heavy-tailed frequency distribution. The spatial maps of the resting state networks (RSNs) obtained via ICA analysis are shown in Fig. 1 and Fig. 2. Order of components is non informative. The computed components account for 56.39% of the data’s variance. An overview of the cortical coverage, as well as the overlap of the components can be found in supplementary Fig. 3. The resulting 30 RSNs are described below following anatomical classification from the Saleem and Logothetis atlas (Saleem and Logothetis, 2012):

**Fig 1.**
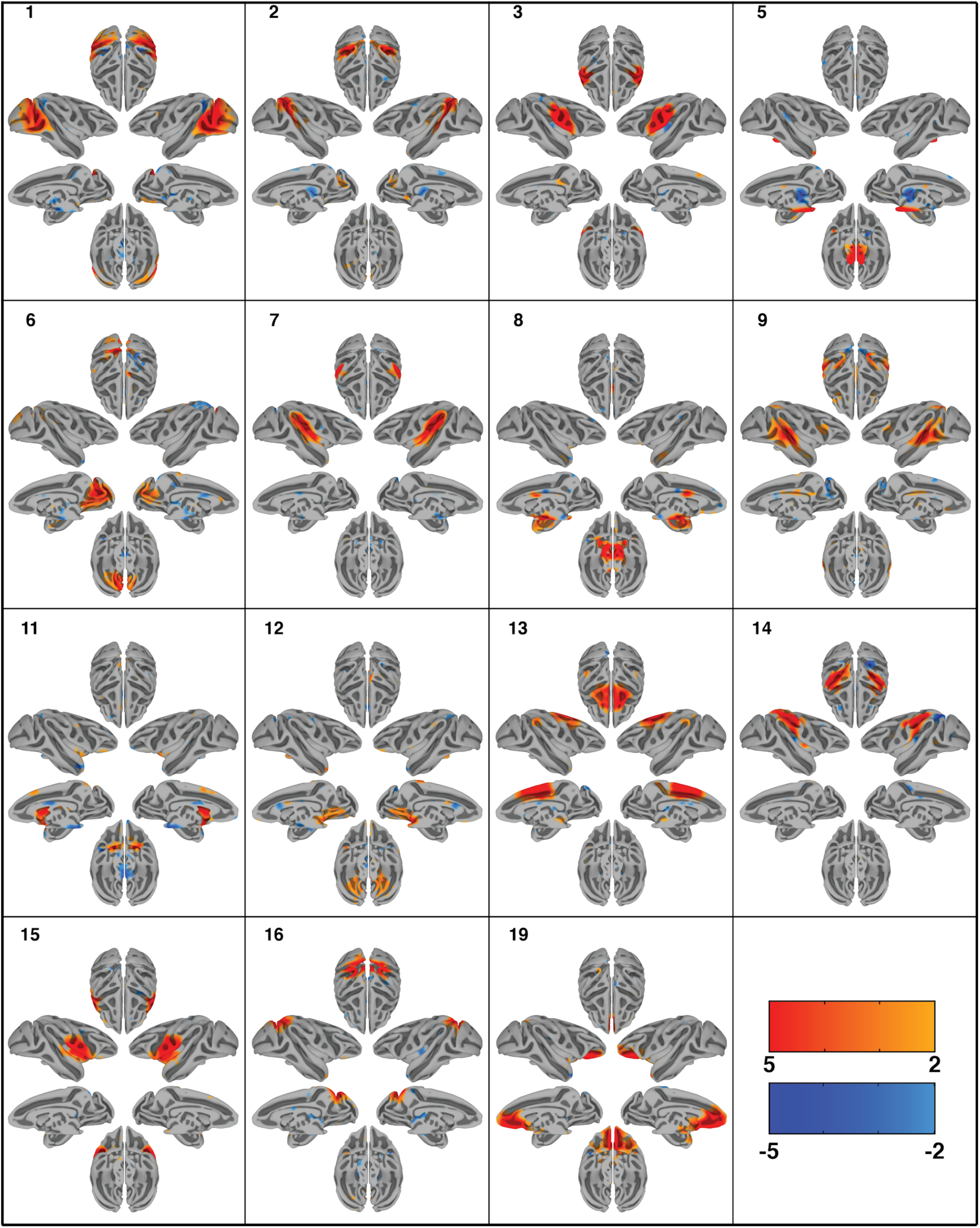
Cortical surface representation of the first 15 resting state networks (RSNs) identified using group independent component analysis (GICA) in 6 macaque monkeys. Overlayed color maps represent thresholded z-scores. Individual subjects were normalized to the NMT template. Each component shows the medial and lateral view of each hemisphere independently as well as the dorsal and ventral view of the hemispheres combined.

**Fig 2.**
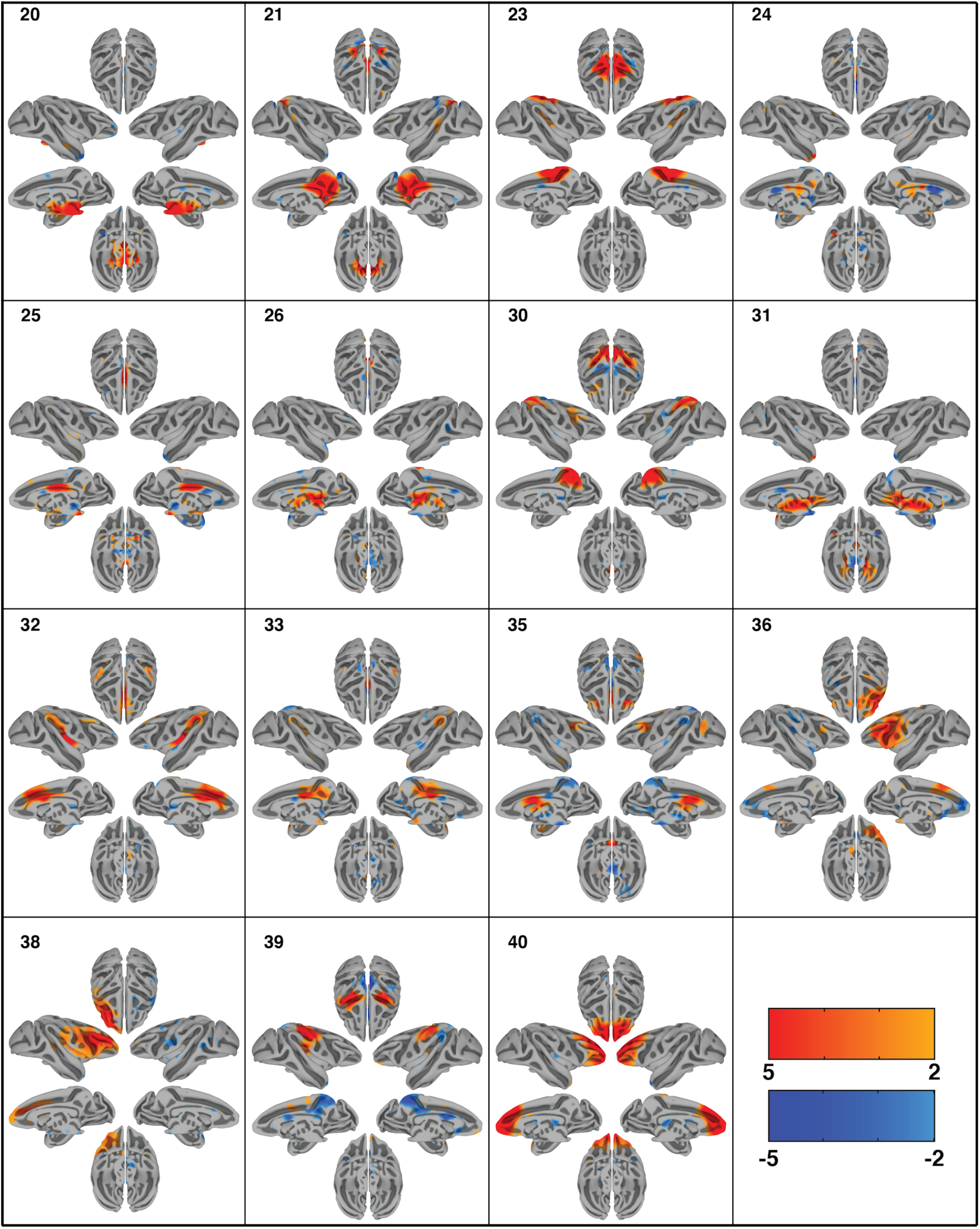
Cortical surface representation of the second 15 resting state networks (RSNs) identified using group independent component analysis (GICA) in 6 macaque monkeys. Overlayed color maps represent thresholded z-scores. Individual subjects were normalized to the NMT template. Each component shows the medial and lateral view of each hemisphere independently as well as the dorsal and ventral view of the hemispheres combined.

- *Network 1 (lateral occipital):* This RSN is mainly in the lateral occipital cortex. It covers portions of early visual areas V1, V2, and V4. The network also includes anticorrelations to regions within the intraparietal sulcus.
- *Network 2 (dorsal superior temporal sulcus):* This RSN includes the areas surrounding the dorsal portion of the STS, including areas 7a, FST, MT, and MST.
- *Network 3 (somatosensory):* This RSN is primarily composed of somatosensory cortical regions, extending somewhat into the primary motor cortex as well. Anticorrelations with insula and lateral intraparietal area are observed.
- *Network 5 (pulvinar-pons-cerebellum):* This RSN is primarily subcortical and includes portions of the dorsal thalamus, particularly the medial and lateral pulvinar, much of the pons, and part of the cerebellum.
- *Network 6 (medial occipital):* This RSN covers medial areas of the occipital lobe. It is primarily composed of area V2, but extends into aspects of areas V6 and, more rostrally, 7m.
- *Network 7 (lateral and cingulate sulci):* This RSN follows the lateral sulcus closely, and thus covers primary auditory cortex, some auditory association areas, area 7op, and large portions of the insula. Although poorly visible on the surface, this network has substantial connectivity with cingulate areas 23c and 24c.
- *Network 8 (ventromedial subcortex):* This RSN covers a large swath of the ventromedial subcortical surface of the brain, including amygdala, basal forebrain, and hippocampus. Cortically, correlations in anterior cingulate cortex 24a/b as well as anticorrelations in nearby 23a/b are observed.
- *Network 9 (middle STS-dlPFC-cingulate):* This RSN covers the middle portions (dorsal-ventrally) of the superior temporal sulcus. This covers many areas associated with multisensory processing, including TPO, TAa, TEO, PGa, and some of the insula. The network also includes portions of the anterior and posterior cingulate gyrus, the outer edges of the intraparietal sulcus, and the dorsolateral prefrontal cortex (specifically areas 46 and 8a). Anticorrelations are seen with visual areas V2 and V6.
- *Network 11 (striatal-medial cortical):* This RSN covers the entire striatum, as well as some medial cortical areas, including precuneus, V6, and dorsomedial frontal areas 6M and 8M. Additional connectivity to orbitofrontal cortex (area 13) and anticorrelations with frontal pole and midbrain are present.
- *Network 12 (cerebellar-frontal pole):* This RSN covers the cerebellum and ventromedial frontal pole.
- *Network 13 (dorsomedial cortex):* This RSN covers a large portion of the dorsomedial cortex, including areas of the frontal lobe (medial areas 9, 8, 6, 4, and 24) and parietal lobe (somatosensory cortices, precuneus, and are 23).
- *Network 14 (intraparietal sulcus):* This RSN covers much of the intraparietal sulcus, including LIP, VIP, and MIP. Rostrally, it extends into the insula.
- *Network 15 (insula):* This RSN is mostly situated in the insula and adjacent operculum. Dorsally, it bleeds into the somatosensory and premotor cortices; ventrally and rostrally, it bleeds into the ventrolateral prefrontal cortex.
- Network 16 (caudal dorsomedial cortex): This RSN is in the dorsal parietal and occipital lobes. It includes portions of V1, V3, V4, V6, precuneus, Opt, LIP, and PEa.
- *Network 19 (orbitofrontal cortex):* This RSN covers nearly all of the orbitofrontal cortex, spanning from orbitofrontal regions of the frontal pole to ventral insula regions of the orbitofrontal cortex. The network extends somewhat dorsally into the medial prefrontal cortex (mainly area 32).
- *Network 20 (midbrain):* This RSN is centered in the midbrain, but extends into portions of the thalamus, hypothalamus, cerebellum, basal forebrain, and pons.
- *Network 21 (posteromedial cortex):* This RSN covers a large portion of the posteromedial cortex, including posterior cingulate areas 23 and 31, parietal area precuneus, and, caudally, some of V6 and V2. There are smaller lateral regions in this network as well: portions of TPO, 7a, and LIP are present. Finally, much of the intraparietal sulcus shows anticorrelations to this RSN.
- *Network 23 (sensorimotor):* This RSN is situated centrally and dorsomedially, spanning the border between the frontal lobe and parietal lobe. Thus, it includes primary motor cortex, primary somatosensory cortex, supplementary motor area, premotor cortex, anterior and posterior cingulate cortices, and PE.
- *Network 24 (lower brainstem)*: This RSN is situated in the pons and medulla.
- *Network 25 (middle-caudal cingulate):* This RSN is mainly situated in the caudal anterior cingulate cortex (area 24’, sometimes called the midcingulate cortex) and posterior cingulate cortex (areas 23 and 31).
- *Network 26 (dorsal thalamus):* This RSN is situated in the dorsal portion of the thalamus.
- *Network 30 (dorsal parietal cortex):* This RSN is located in the dorsal parietal cortex, and covers the intraparietal sulcus, including areas LIPv, LIPd, area 5 and MIP. It extends towards the medial wall, covering areas V6, precuneus, and parts of the posterior cingulate cortex.
- *Network 31 (ventral thalamus):* This RSN is located in the ventral thalamus, with some extension into the midbrain.
- *Network 32 (anterior cingulate-insula):* This RSN covers the entire anterior cingulate cortex (area 24), much of the insula, and a small portion of the lateral parietal cortex (7op and PFG).
- *Network 33 (posterior cingulate-lateral parietal cortices):* This RSN covers the posterior cingulate cortex (areas 23 and 31) as well as lateral parietal areas 7op and Tpt. There is also a small area of correlation in the temporal pole.
- *Network 35 (dorsolateral-cingulate-thalamus):* This RSN includes areas in the dorsolateral prefrontal cortex such as area 8 as well as the anterior cingulate cortex area 24 and some rostral thalamus. Anticorrelations to posterior cingulate cortex and precuneus regions are observed.
- *Network 36 (lateral prefrontal right):* This RSN covers the right lateral prefrontal cortex around the principle and arcuate sulcus. Anticorrelations with the medial prefrontal cortex, such as area 9 and 25, as well as part of the anterior cingulate cortex are observed.
- *Network 38 (lateral prefrontal left):* This RSN covers the lateral left prefrontal cortex around the principle and arcuate sulcus.
- *Network 39 (sensorimotor 2):* This RSN covers both sides of the central sulcus, and thus involves both motor and somatosensory cortices. The network also includes anticorrelations of posterior cingulate cortex and more caudal somatosensory areas.
- *Network 40 (frontal pole):* This RSN is mainly in the frontal pole. Caudally, it extends into the dorsal prefrontal cortex and anterior cingulate cortex.

### Individual resting state networks

Since group analysis can omit individual anatomical and functional details, we present all individual ICA results in supplementary Figs. 4-9. Figure 3 shows the consistency of our single subject independent component results in all subjects for four representative components. Note the consistency in spatial location at the same threshold. This finding demonstrates the consistency and data quality of our imaging approach and shows that our group level findings are highly representative of the individual monkey.

**Fig 3.**
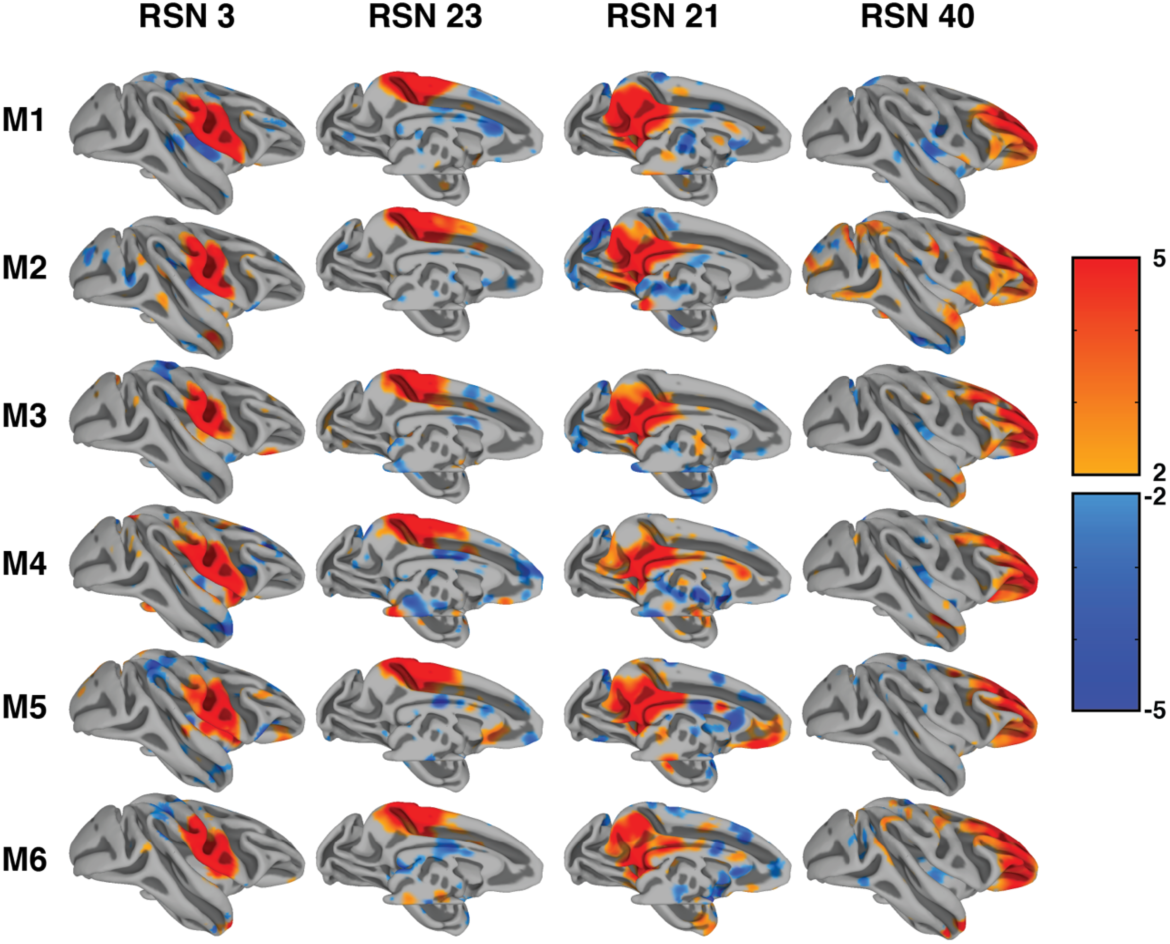
Consistency of single subject resting state networks. The figure demonstrates the same four resting state components in all six monkeys. Thresholds were kept consistent with group analysis results.

Projection to surface space can additionally obscure results and is generally ill-suited for use in individual animal physiological use. Fig 4. shows 7 of the 30 components displayed from one representative single subject volume space projected onto the NMT template. The results demonstrate the close correspondence of the components with anatomical and functional landmarks as well as specificity to both cortical and subcortical regions.

**Fig 4.**
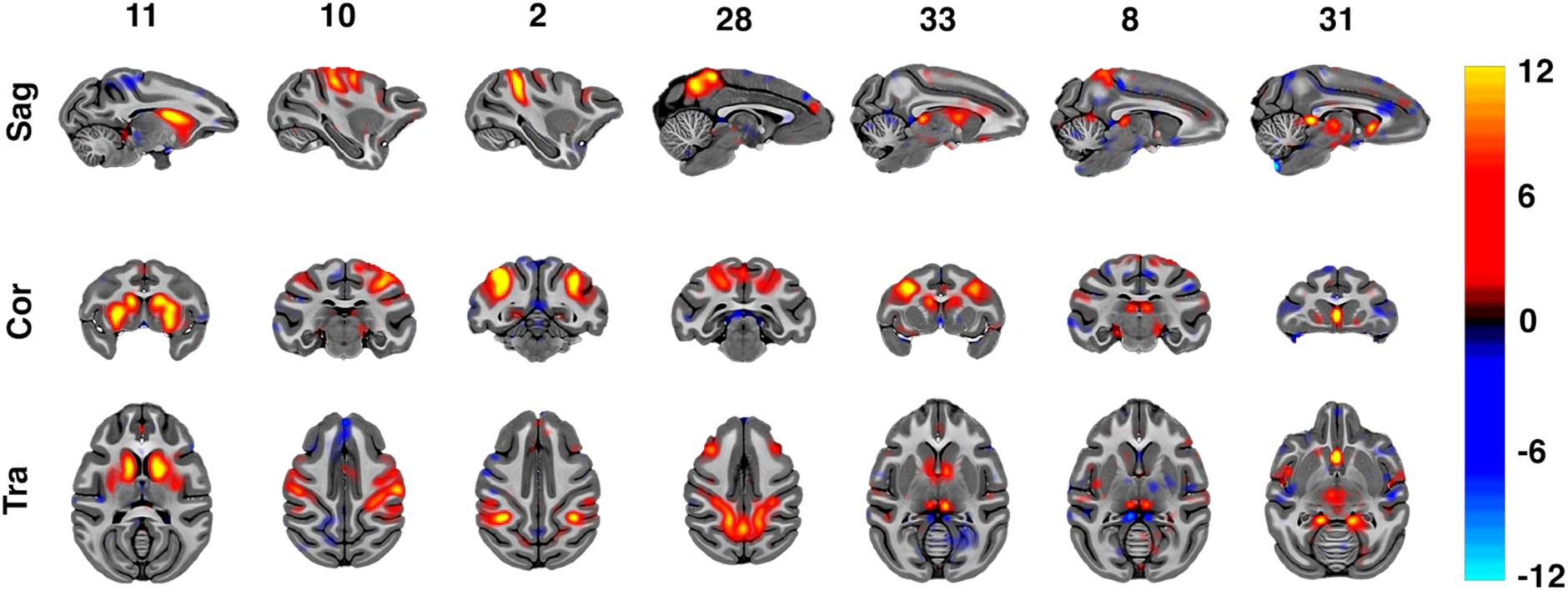
Example resting state networks (RSNs) of one macaque following single-subject independent component analysis (ICA). Images are normalized to the NMT space and overlaid onto the NMT-template. Overlaid color maps represent thresholded z-scores. Arbitrary slices were selected to demonstrate the diversity of components from subcortical to cortical components as well as the detailed anatomical correspondence.

### Independent component network connectivity

Data driven hierarchical clustering of the independent component time courses revealed 9 clusters of components, with strong positive functional connectivity within cluster and between adjacent clusters, and strong negative functional connectivity between non-adjacent clusters. Even though clustering was not based on spatial information, the resulting adjacencies reveal anatomical neighboring relationships. Clusters fall into the following categories: 1. Subcortical components, 2. Prefrontal components, 3. Visual components, 4. Parietal components, 5. Superior-temporal components and 6. Motor components.

## Discussion

Here we present the first demonstration of connectome style quality resting state fMRI data in the macaque at ultra-high magnetic fields. The delineated RSNs closely resemble the quality and findings of connectomes from large human datasets, providing a first detailed map of the rsfMRI ‘macaque connectome’ as a basis for future interventional work.

Resting state functional connectivity patterns are an extremely important tool used in current neuroscientific research into the origin of mental disease states (Bai et al., 2011; Lawrie et al., 2002; Lowe et al., 2002; Lustig et al., 2003; Quigley et al., 2001; Xie et al., 2012). Yet, much of this technique’s physiological basis is not well understood. Since rhesus macaques offer an extremely promising translational model (Consortium et al., 2020; Hutchison et al., 2011; Hutchison and Everling, 2012; Milham et al., 2018), a detailed map of the rsfMRI ‘macaque connectome’ is needed to build the basis for future interventional work. While human rsfMRI studies have made significant advances in the ability to acquire highly sensitive and informative images of functional connectivity (Essen et al., 2012; Glasser et al., 2016; Harms et al., 2018), including at relatively high spatial and temporal resolutions (Moeller et al., 2010; Vu et al., 2016), similar non-invasive neuroimaging advances in the macaque have not been as widespread or productive.

Several barriers have previously prevented such translations from being successful. Since invasive work, such as single unit recording, anatomical dissection, or chemo- and optogenetic manipulations in the macaque is usually done on the individual subject level at high spatial and temporal scales, extremely robust within individual rsfMRI maps are needed. In vivo experiments using ultra-high magnetic fields facilitate such high spatial-temporal resolutions and overall sensitivity across the entire brain. Next, high channel count coils with sufficient parallel imaging performance, specifically designed for macaque setups, are also essential to achieving the translational sensitivity and spatial-temporal resolutions. Further, even though human studies are typically completed within a couple of hours and macaque studies have the luxury of extending well beyond this to increase overall sensitivity, extending session duration can still be problematic in terms of physiological (and therefore fMRI) stability. For this reason, shorter scans with higher SNR are still desirable.

Another approach to achieving sensitivity with shorter scan durations is the use of contrast agents, which even makes lower field studies feasible. However, the use of contrast agents has significant limitations as well, such as: 1) It does not measure intrinsic BOLD signals (which is what the vast majority of human studies use), 2) studying temporal dynamics or paradigms other than block designs is much more difficult, and 3) there are risks associated with its use and it is hence not used in any human studies. Next, while implementation on a human-based scanner is not absolutely necessary, it does provide a convenient avenue by which imaging methods and sequences routinely used in the human can be trivially applied to the macaques. Finally, while obtaining high resolution functional images of the alert monkey would be more ideal than the low anesthesia strategy we employed here, this is still an enormous technical challenge that we and others are working towards. Despite this, our data demonstrated superb functional contrast to noise and the setup described here could be used for other applications, such as task fMRI.

Here, building on previous work (Hutchison et al., 2011), we report the first whole brain ultra-high resolution (0.75 mm^3^ isotropic at ∼1.2s TR) comprehensive group independent component analysis in macaques at 10.5 T. Our approach identified 30 reproducible resting state networks covering the entire macaque cortex as well as, for the first time, subcortical contributions throughout the thalamus, brainstem and cerebellum. The RSNs we were able to identify closely resemble higher model order RSNs identified in large sample sizes of human experiments (Abou-Elseoud et al., 2009; Kiviniemi et al., 2009; Li et al., 2007; Smith, 2012; Smith et al., 2013, 2012, 2009). In these findings, high model order RSNs are a tradeoff between the number of components extracted and the detectability of unique networks observed. Extremely high model order fraction larger networks but tend not to add additional unobserved region involvement. Our networks represent an intermediary between extremely high model orders (Moeller et al., 2009) and more conservative approaches (Hutchison et al., 2011), covering nearly the entire brain (see Fig. 6 and supplementary Fig. 3) while uncovering networks that include long range functional connections. Using this approach, we were able to demonstrate remarkable reproducibility of the RSN decomposition even in the individual animal without the use of contrast agents. This achievement is of utmost importance when trying to understand and model the relationship between real anatomical neural connections only obtainable invasively and functional connectivity based on neuroimaging. Our RSN findings reveal very similar network organization between humans and macaques at the mesoscopic level (see Fig. 6 for a qualitative comparison). We observe RSNs in all of the networks that are commonly identified in human experiments (Beckmann et al., 2005; Jafri et al., 2008; Smith et al., 2013, 2009). For example, we identify multiple networks that resemble the commonly described fronto-temporal, default-mode, fronto-parietal, cerebellar, visual, somato-motor, intraparietal, lateral prefrontal, basal ganglia, insular, cingular networks (Fig. 6). This is the first time such a comprehensive and high quality has been obtained in macaques and the world’s first such dataset at 10.5 Tesla.

**Fig 5.**
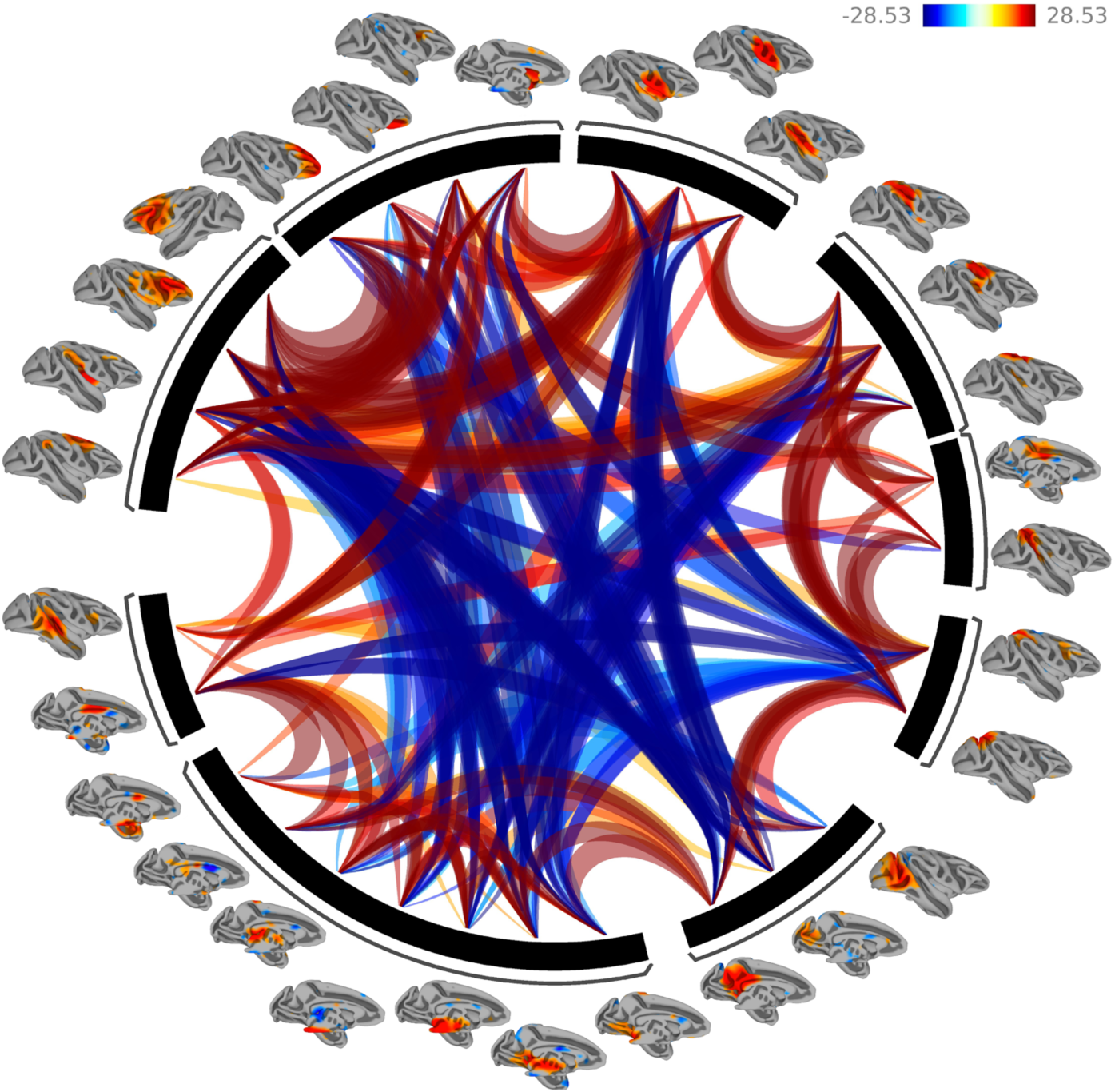
Functional Network Connectivity of the 30 extracted Independent components. Lines portray significant connections between the networks and color indicates positive (red) and negative (blue) functional connectivity. Networks were clustered using hierarchical clustering. Results demonstrate strong positive within cluster network connections and primarily negative between cluster connections.

**Fig 6.**
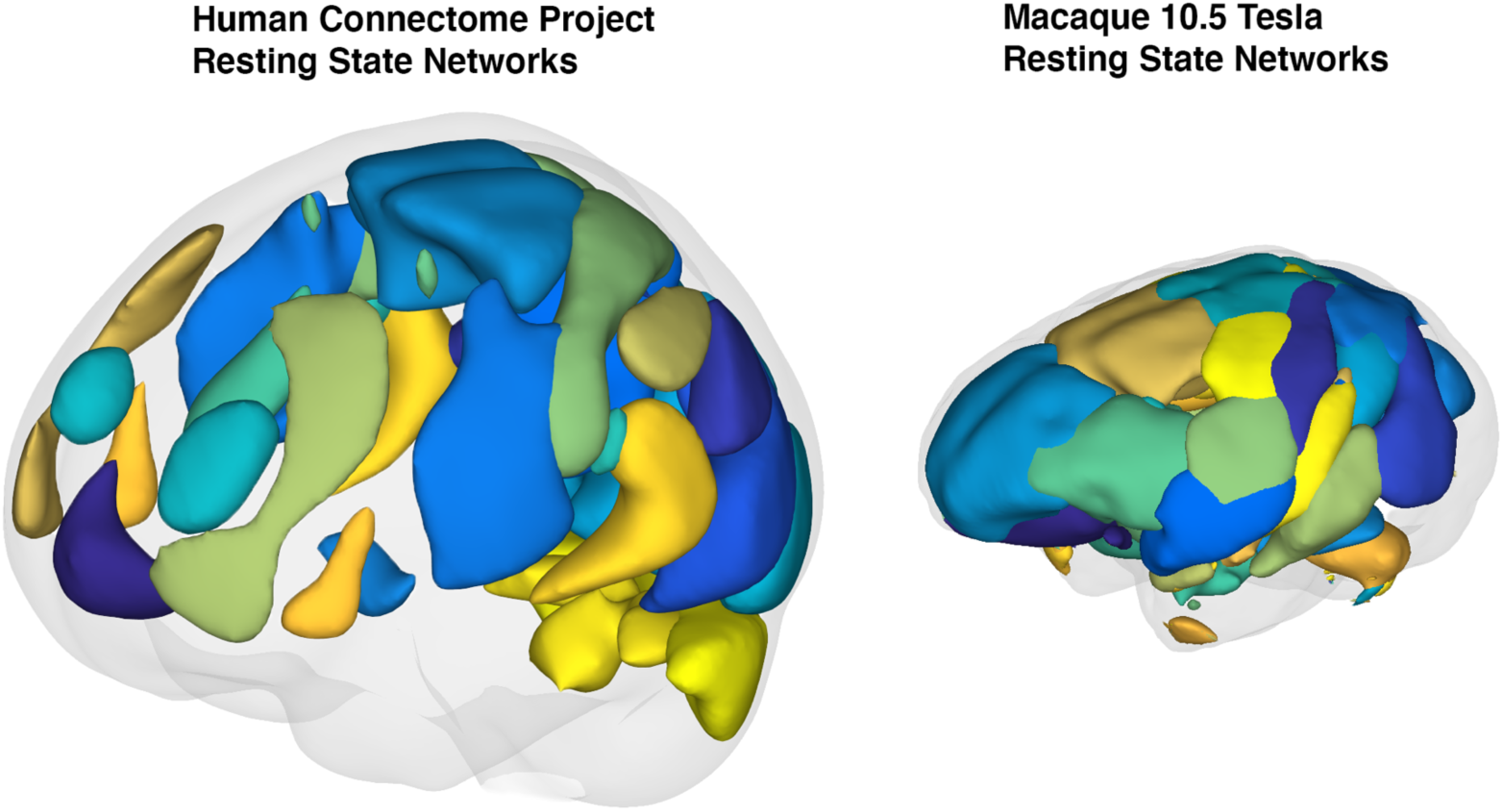
Qualitative comparison between resting state networks identified from the Human Connectome Project database (supplied by the CONN toolbox and Connectome Workbench) and our six subject macaque group resting state networks. Individual regions of interest were smoothed using a 10 mm spherical kernel interpolated in 3D. Gray shading represents a reconstruction of the estimated pial surface.

Previous work found 20 stimulus evoked components (Moeller et al., 2009) using contrast agents or 11 high quality intrinsic components (Hutchison et al., 2011). Extending this work, we were able to resolve previously not observed (Hutchison et al., 2011; Moeller et al., 2009; Vincent et al., 2009, 2007) lateralized dorsolateral prefrontal components (RSN 36,38) that have readily been described in the human literature and attributed to language, memory and cognitive attentional processes (Beckmann et al., 2005; Jafri et al., 2008; Smith et al., 2009). Our lateralized network does not include the parietal contribution present in the human literature, however. Another set of components that have previously remained elusive (Vincent et al., 2009, 2007) in the macaques are the dorso medial and orbital contributions of the prefrontal cortex. In our findings (RSN 19, 40) these were readily present. It is difficult to assess why these findings have previously not been reported, but SNR limitations as well as susceptibility artefacts close to the orbits and sinuses have likely contributed to the problem. One difference in our findings with the human literature is the absence of a large component covering the primary visual cortex’s caudal extent. Our visual cortex related RSNs (RSN 1,6) cover both medial and lateral portions of V1, V2 but miss the occipital pole. This is likely the result of our coil having the lowest sensitivity in visual cortex due to the macaques sphinx position. Another difference in our findings compared to previous macaque rsfMRI studies (Hutchison et al., 2011; Moeller et al., 2009; Vincent et al., 2009, 2007) but in agreement with high model order RSN studies (Abou-Elseoud et al., 2009; Kiviniemi et al., 2009) as well as clustering approaches (Leech et al., 2012, 2011), is the fractioning of RSNs that are associated with the default mode network. In our findings multiple RSN (21, 26, 30, 31, 33, 39) show at least partial overlap with posteromedial cortex. Another finding that has previously not been reported is the detailed delineation of the cingulate cortex that can be observed in our RSNs. RSNs (5,8,9,11,13,21,24,25,32,33,35,39) all exhibit involvement of the cingulate cortex partitioning the anatomical areas 23a,b,c and 24a,b,c. While some of these networks demonstrate overlap in their cingulate cortex involvement, unique partitioning is directly observable in others (RSNs 5,24,25,26,35). Again, it is unclear why these effects were not previously found but inhomogeneous excitation from local transmitters or low SNR in medial regions further away from the scalp (especially true if headposts are used) could contribute to this omission.

To address one of our motivations, the robust identification of RSNs within the individual subject, single subject ICA using the same model order of 40 were performed and assessed for reproducibility and compared to the group results. Compared to previous studies (Hutchison et al., 2011) we found remarkable reproducibility for most of the 30 resulting RSNs. First, using the same thresholds as in the group-ICA resulted in more widespread diffuse activity, yet the pattern and anatomical correspondence of the main group-ICA findings were largely preserved in the individual. It is difficult to compare our current approach to previous findings since we used diffeomorphic transformations for normalization that likely resulted in better anatomical correspondence as well as acquired significantly more data in the individual subject.

Taken together, we have achieved a highly detailed, individually robust delineation of RSNs in the macaque that closely resembles the quality and findings of connectomes from large human datasets, further demonstrating the benefits of using ultra-high magnetic fields in vivo. One countering recent argument and effort (Consortium et al., 2020; Milham et al., 2018) that has seen high translational success has been the cross laboratory pooling of macaque fMRI datasets. We argue that our rationale of advocating ultra-high field macaque neuroimaging does not negate or change the need for such databases, but rather aims at providing data and solutions for a different problem. The big data consortium approach allows for the study of general principles derived from rsfMRI data in macaques such as physiological states (Xu et al., 2019) (anesthetized vs. awake) or macroscale area organizations (Xu et al., 2018) within the species. These are incredibly important efforts that can also serve an important translational role. What these approaches cannot provide, and where we believe ultra-high field comes into play, is measuring the effects of within subject manipulation using invasive tools. If we are interested in figuring out how structural connections and their electrical and chemical changes enable functional connectivity in rsfMRI and behavior, disease or developmental processes, resting state neuroimaging at ultra-high magnetic fields coupled with within-subject invasive manipulations will pave the way.

## Conclusions

To our knowledge, this is the first demonstration of a macaque connectome (or HCP) style resting state dataset, which has been available for human applications for around 10 years. While previous macaque resting state studies have demonstrated typical RSNs, none have achieved whole brain sub-millimeter acquisitions with an approximate 1 sec temporal resolution, let alone do it using only intrinsic BOLD signals and with a scan duration of around an hour. Data sets such as the current one allow for more straightforward translations of macaque data to the human for the purposes of understanding brain connectivity in individuals and how it is altered in disease.

## Acknowledgements

We thank Steve Jungst for continuing support with our coils and hardware setup. We thank Hannah Lee, Jen Holmberg, Adriana Cushnie, Tanya Casta, and Megan Monko for support with animal care and data acquisition. We thank Research Animal Resources at UMN, especially Whitney McGee and Anne Merley, for helping us implement new and improved anesthesia protocols.

## Supplementary Materials

**Supplementary Figure 1.**
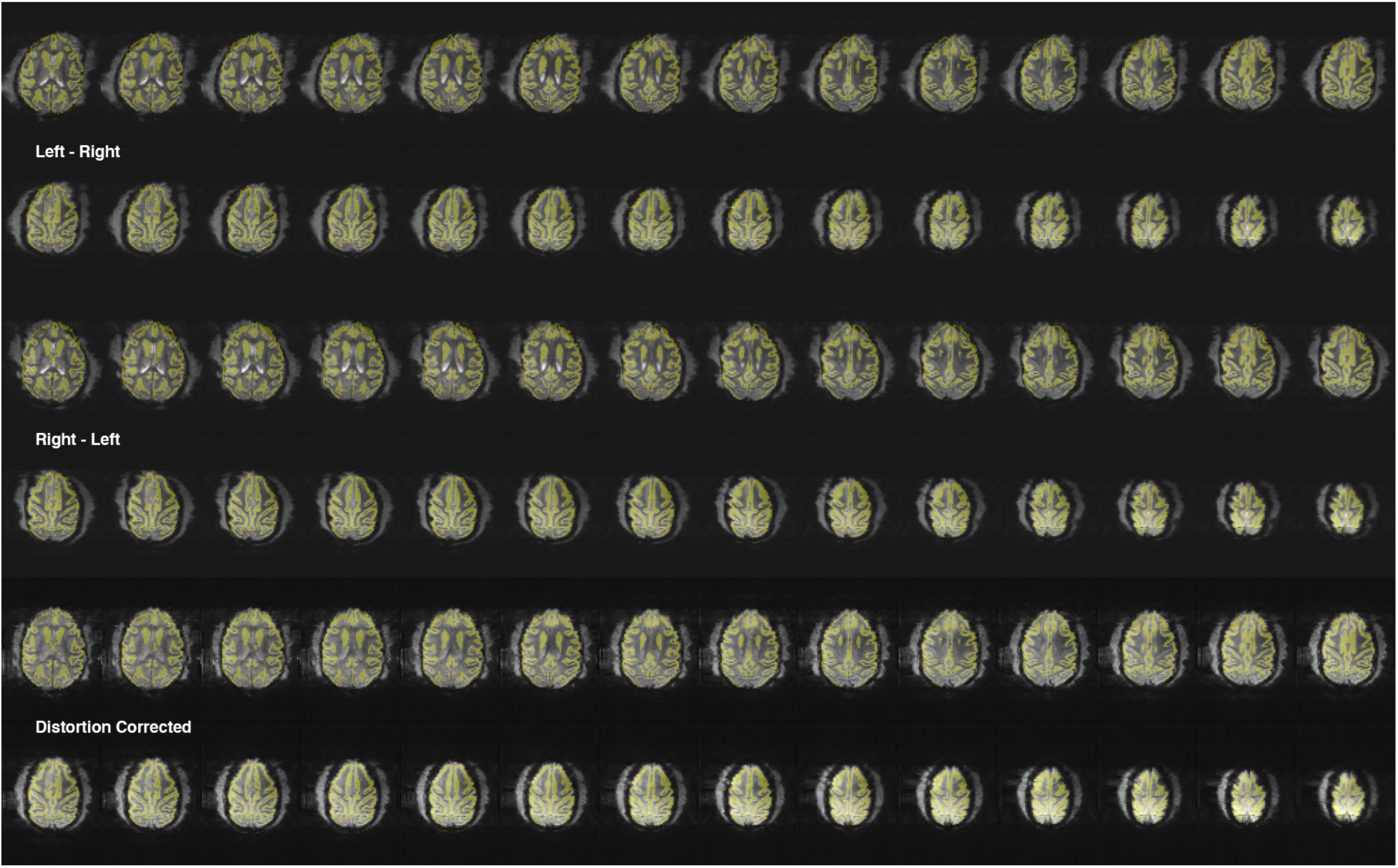
EPI distortion correction results. Selection of transversal slices of one macaques raw gradient echo 2D EPI data for both phase encoding directions (LR-RL, top 4 rows) as well as the EPI distortion corrected image used in our analysis (bottom 2 rows). Gray matter segmentation results from the individual anatomical T1 image are overlaid in opaque yellow without registration (scanner position only). The distortions we experience are small to start with and the distortion correction provides very good correction leading to near perfect alignment without the use of any registration procedures.

**Supplementary Figure 2.**
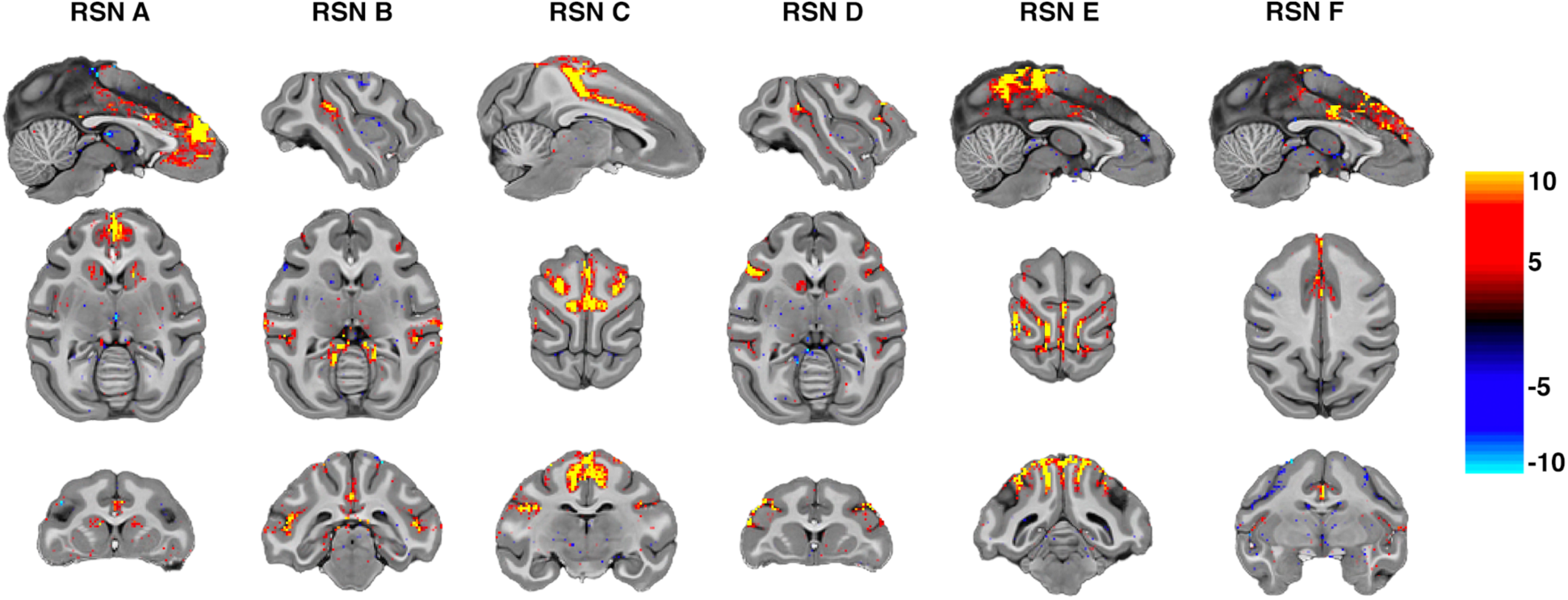
Independent components of unsmoothed data in individual subject space. Displayed are 6 RSNs in one unsmoothed macaque dataset. For this visualization thresholds were set to 4,-4. The results show robust results in the unsmoothed volumes at native resolution demonstrating our high SNR regime. Networks adhere to the cortical ribbon without any registration performed.

**Supplementary Figure 3.**
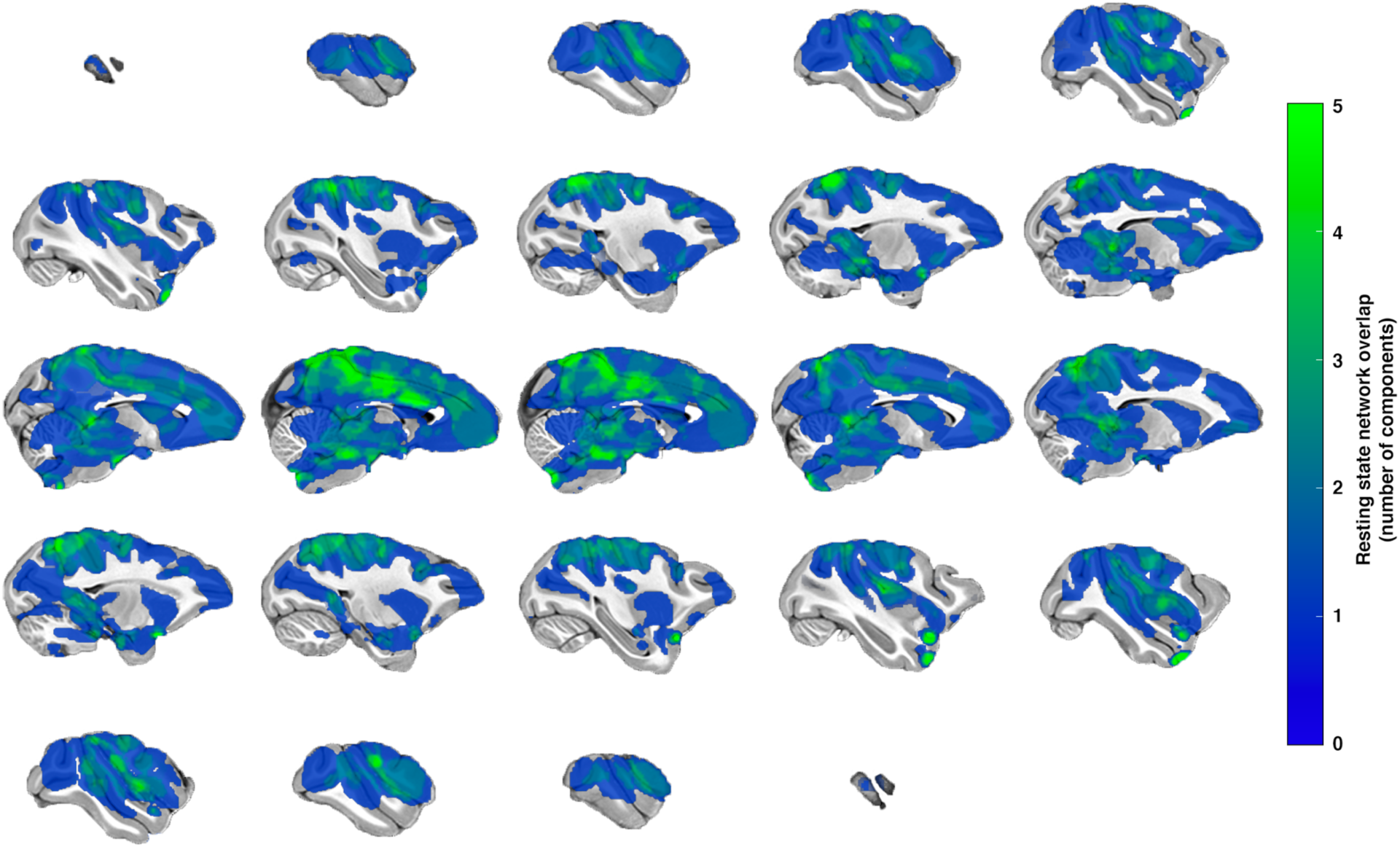
Sagittal slices displaying the overlap of all 30 analysed resting state networks resulting from the Group-ICA. The colorbar indicates the total number of resting state components that have activity (both positive and negative correlations are combined) at the indicated location.

**Supplementary Fig 4.**
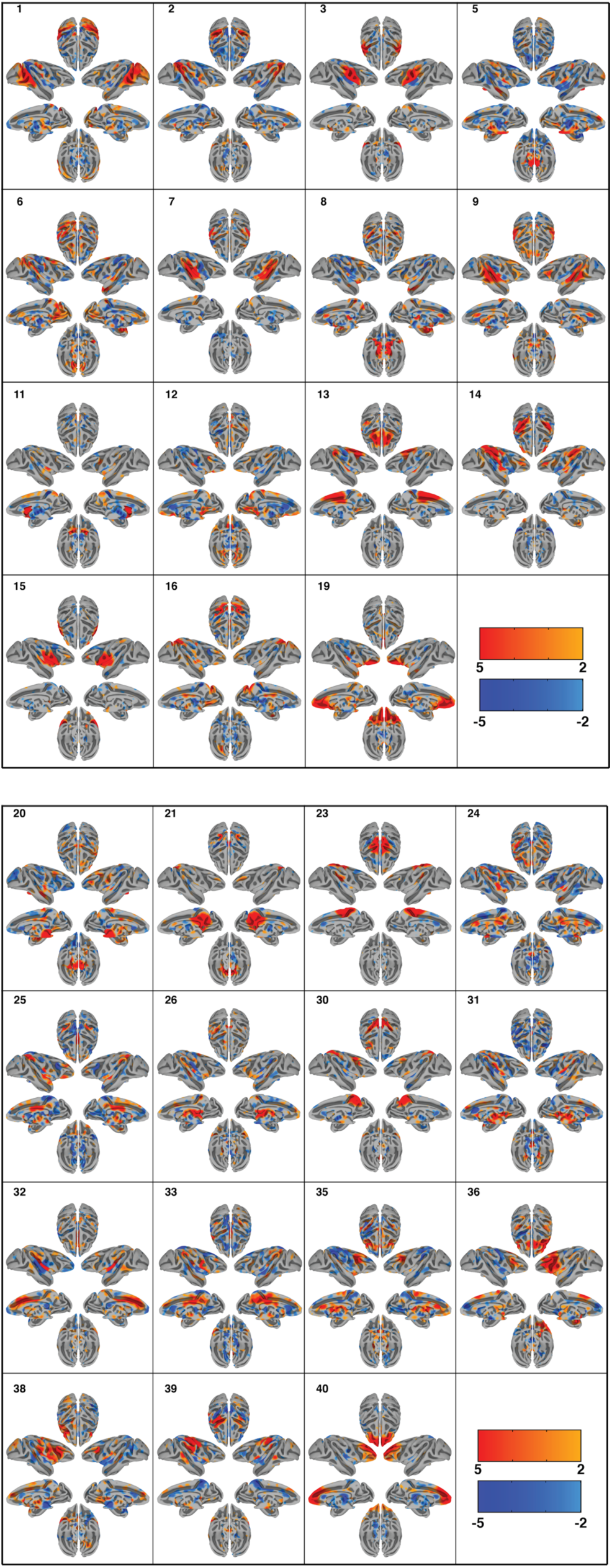
Cortical surface representation of the resting state networks (RSNs) identified using group independent component analysis (GICA) in monkey 1. Overlayed color maps represent thresholded z-scores. Data was normalized to the NMT template. Each component shows the medial and lateral view of each hemisphere independently as well as the dorsal and ventral view of the hemispheres combined.

**Supplementary Fig 5.**
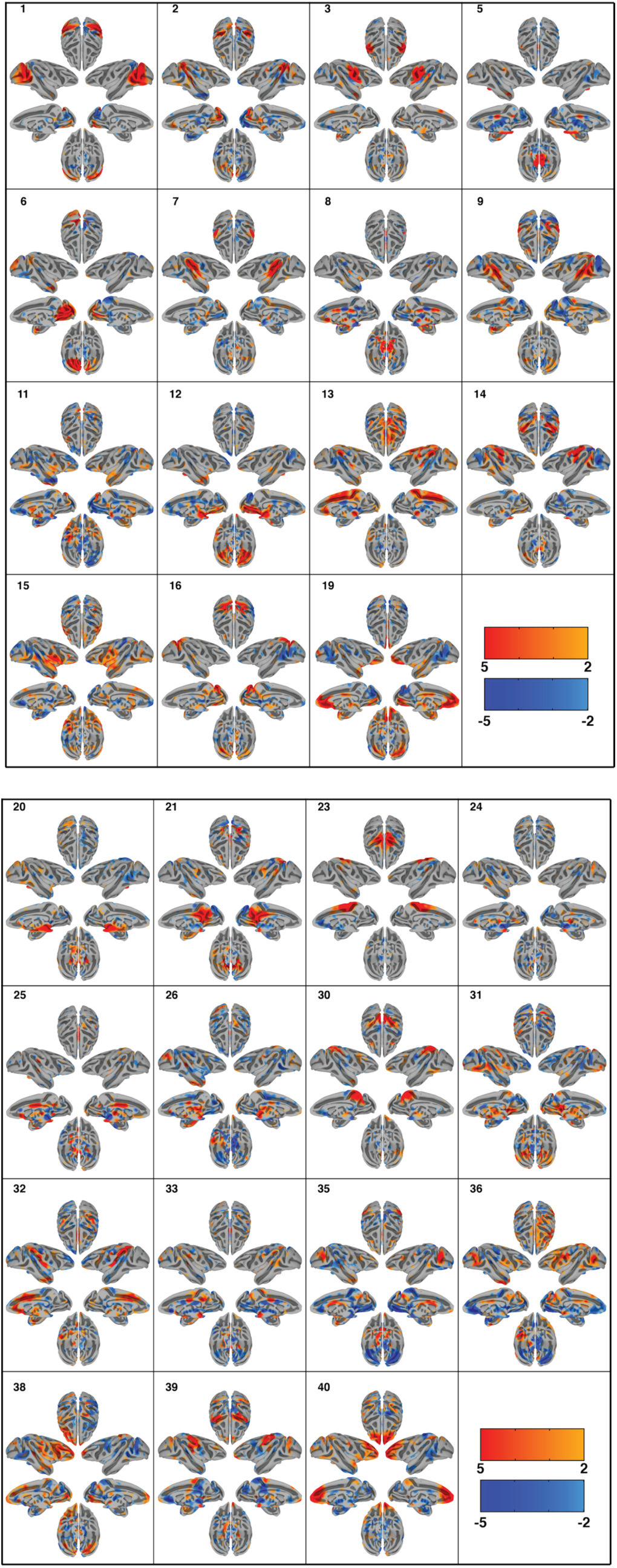
Cortical surface representation of the resting state networks (RSNs) identified using group independent component analysis (GICA) in monkey 2. Overlayed color maps represent thresholded z-scores. Data was normalized to the NMT template. Each component shows the medial and lateral view of each hemisphere independently as well as the dorsal and ventral view of the hemispheres combined.

**Supplementary Fig 6.**
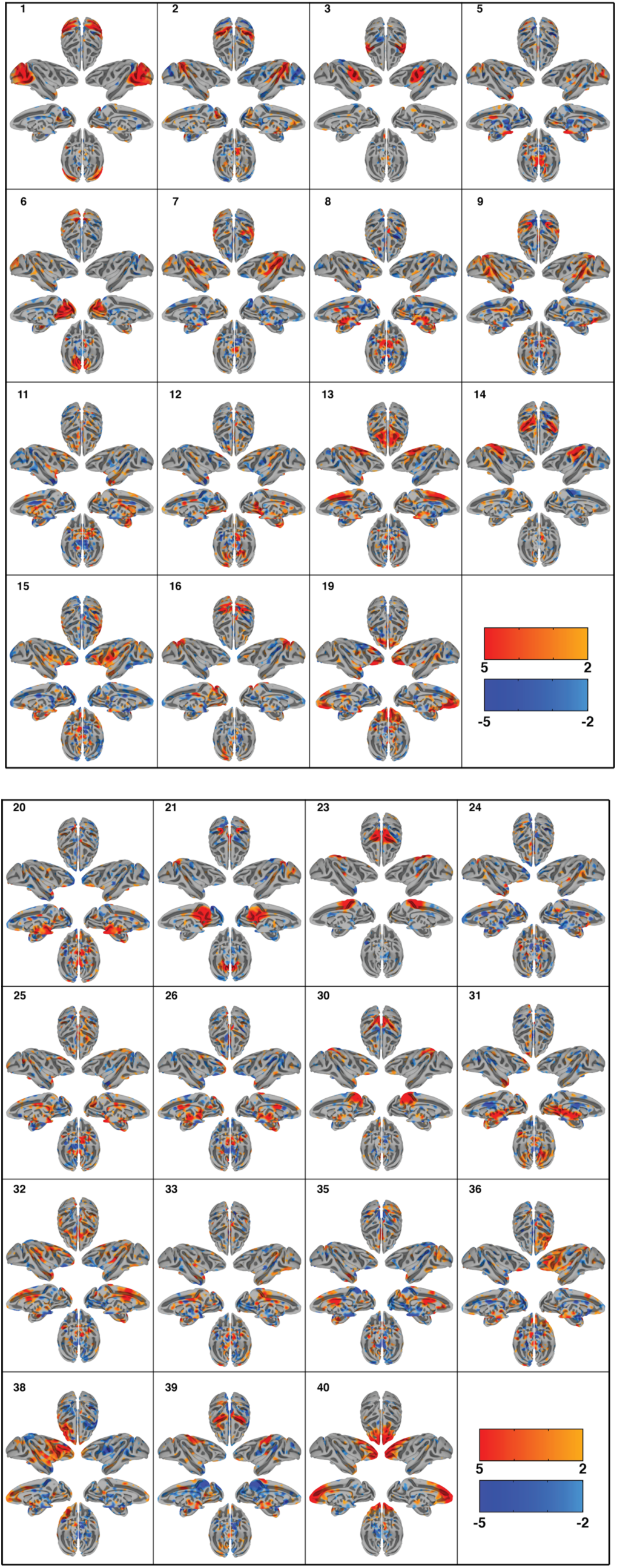
Cortical surface representation of the resting state networks (RSNs) identified using group independent component analysis (GICA) in monkey 3. Overlayed color maps represent thresholded z-scores. Data was normalized to the NMT template. Each component shows the medial and lateral view of each hemisphere independently as well as the dorsal and ventral view of the hemispheres combined.

**Supplementary Fig 7.**
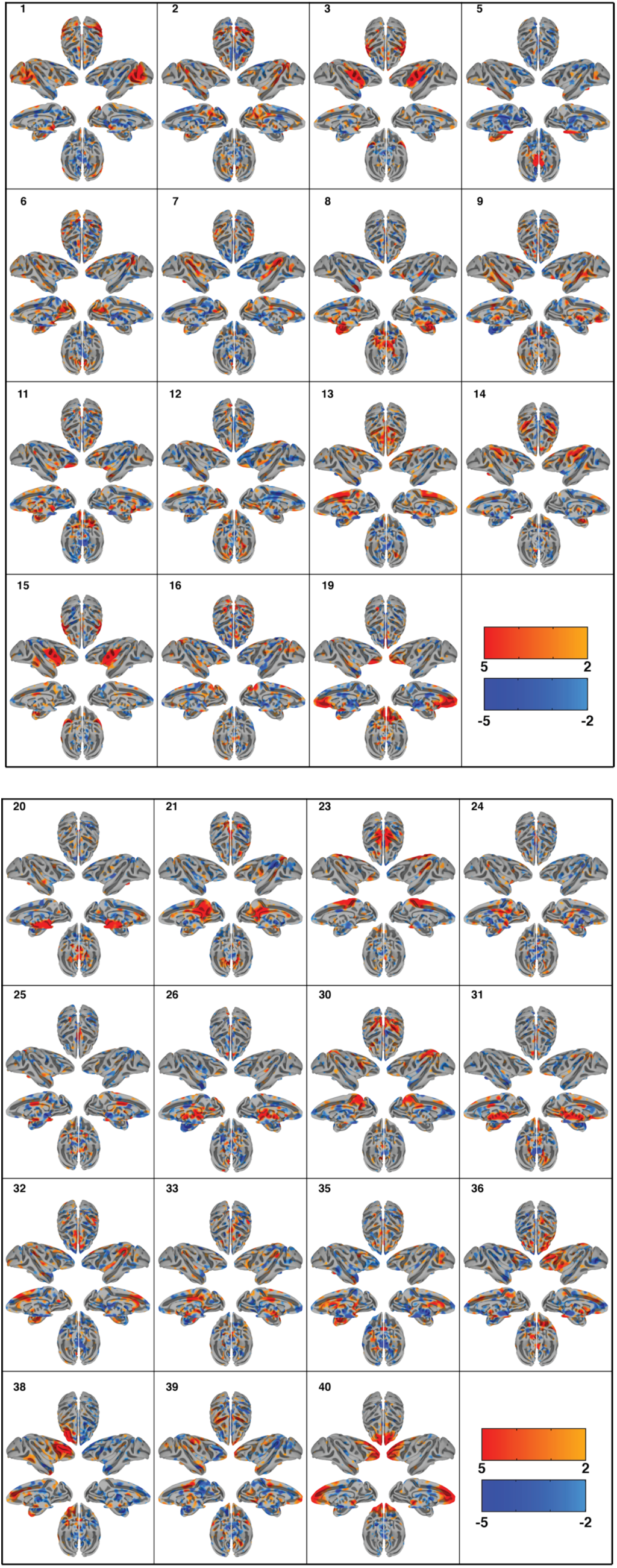
Cortical surface representation of the resting state networks (RSNs) identified using group independent component analysis (GICA) in monkey 4. Overlayed color maps represent thresholded z-scores. Data was normalized to the NMT template. Each component shows the medial and lateral view of each hemisphere independently as well as the dorsal and ventral view of the hemispheres combined.

**Supplementary Fig 8.**
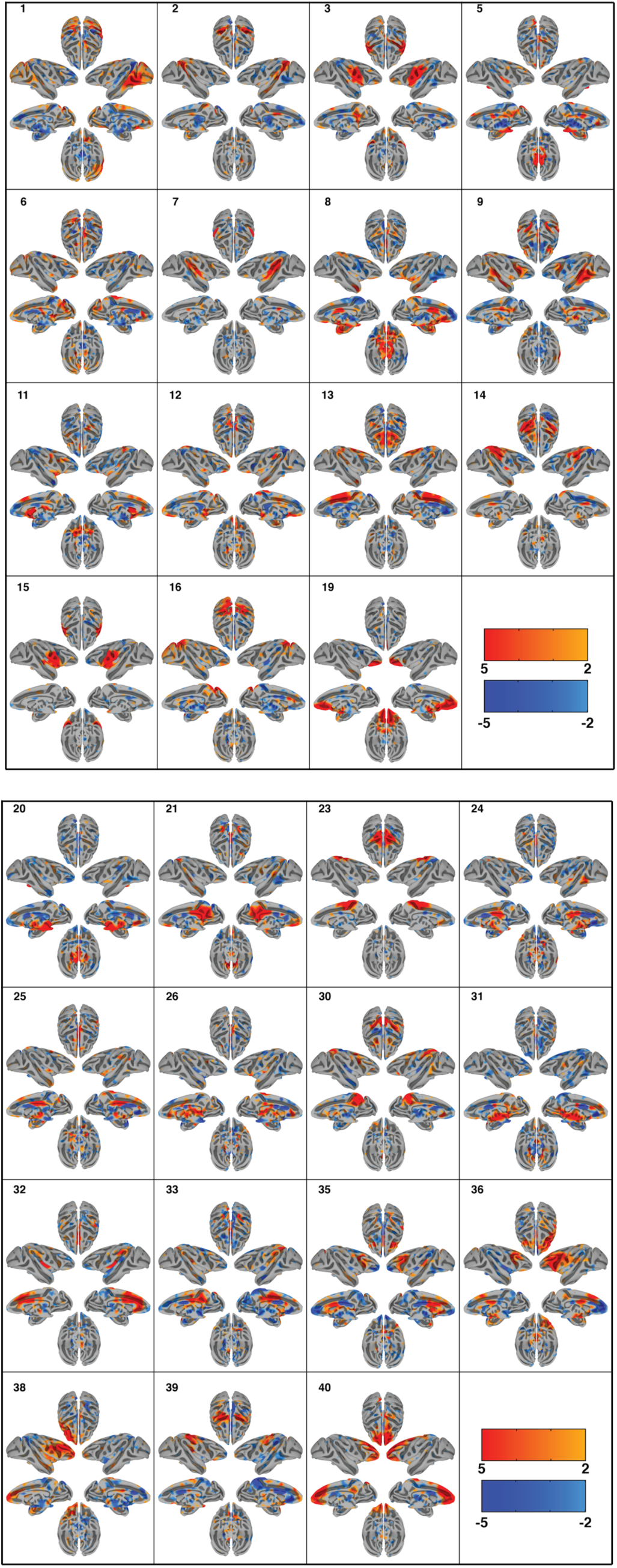
Cortical surface representation of the resting state networks (RSNs) identified using group independent component analysis (GICA) in monkey 5. Overlayed color maps represent thresholded z-scores. Data was normalized to the NMT template. Each component shows the medial and lateral view of each hemisphere independently as well as the dorsal and ventral view of the hemispheres combined.

**Supplementary Fig 9.**
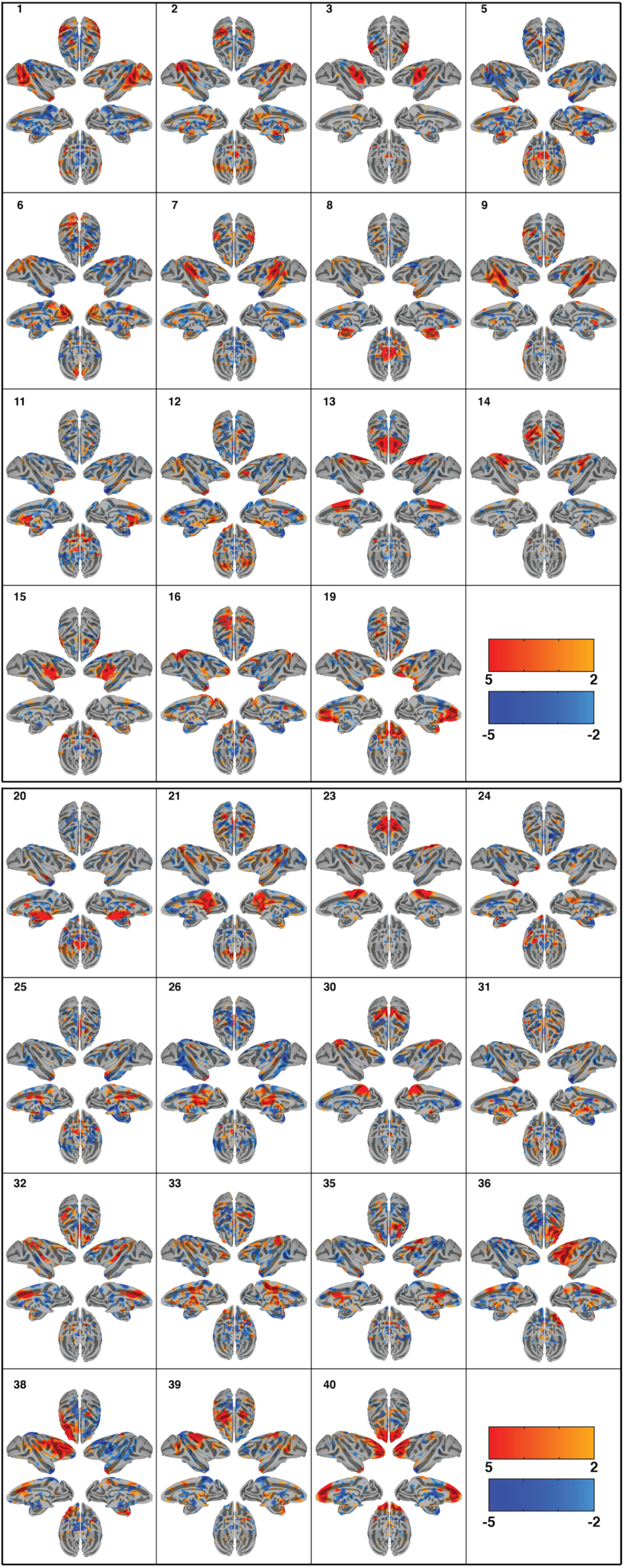
Cortical surface representation of the resting state networks (RSNs) identified using group independent component analysis (GICA) in monkey 6. Overlayed color maps represent thresholded z-scores. Data was normalized to the NMT template. Each component shows the medial and lateral view of each hemisphere independently as well as the dorsal and ventral view of the hemispheres combined.

## Notes

Funding statement This work was supported by NIH grants RF1 MH116978 (to EY), R01 DA038615 (to BYH), U01 EB025144 (to KU), by a P41 EB027061 (to KU, EY, NH, GA, BYH and JZ), R01 MH118257 (to SRH), an NINDS R01 NS081118 and P50 NS098573 Udall center to NH by an award from MNFutures to BYH, from the Digital Technologies Initiative to JZ, and BYH, from the Templeton Foundation to BYH, a Young Investigator Award from the Brain & Behavior Research Foundation to SRH, a Medical Discovery Team on Addiction Pilot Grant to SRH and BYH, and a UMN AIRP award to JZ, BYH, SRH and AZ.

### Competing Interest Statement

The authors have declared no competing interest.

